# Invasive alien predators overturn the spatial-scaling laws of biocomplexity

**DOI:** 10.64898/2026.04.16.718936

**Authors:** Pierre-Gilles Lemasle, Jean-Marc Paillisson, Jean-Marc Roussel, Rémi Lacroix, Paule Lacroix, Gérard Lacroix, Eric Edeline

**Affiliations:** DECOD (Ecosystem Dynamics and Sustainability), INRAE, Institut Agro, Ifremer; Rennes, France; ECOBIO (Ecosystèmes, Biodiversité, Evolution), University of Rennes, CNRS, Rennes, France; Institut du développement et des ressources en informatique scientifique (IDRIS), CNRS, Orsay, France; 9 résidence des Coteaux, 78460 Chevreuse, France; Institut d’écologie et des sciences de l’environnement (iEES Paris), Sorbonne Université, CNRS, UPEC, CNRS, IRD, INRA, Paris, France

**Keywords:** Biological invasions, Biocomplexity-area relationships, Food-web assembly, Top-down ecosystem control, Trophic theory of island biogeography

## Abstract

The theory of island biogeography and its trophic extensions predict that both species richness and food-web complexity should increase with increasing ecosystem surface area. Accordingly, Species-Area Relationships (SARs) and Network-Area Relationships (NARs) are often observed to be positively-sloped, an observation that came to be considered as a law, and on which rest many area-based conservation plans for biodiversity. However, our mechanistic understanding of the driving mechanisms of SAR’s and NAR’s slopes remains limited, undermining our ability to predict how biodiversity will respond to habitat gain or loss. We show in 180 rural ponds sampled across five years that invasive alien predators reversed the SAR and NARs from positive in invader-free ponds, to negative in invaded ponds. Relationship reversal resulted from a higher prevalence of invasive alien predators driving magnified prey extinctions and simplified food webs in larger ponds. The ability of invasive alien predators to reverse SAR and NARs presumably reflected disproportionately high predation rates combined with a low sensitivity to prey extinction conferred by a wide trophic generalism. In a world where virtually all ecosystems face biological invasions, omnipresent invasive alien predators stress the pivotal role played by predation in shaping biocomplexity-area relationships, and highlight a growing need to preserve small ecosystems where invasive alien predators are less prevalent.

## Introduction

The species–area relationship (SAR) describes how ecosystem surface area determines the number of species found within that ecosystem. Empirically, larger ecosystems are often observed to harbor larger numbers of species, resulting in positive-sloped SARs (1, 2). Accordingly, the theory of island biogeography predicts that larger ecosystems have a wider niche space and support larger populations, such that species extinction rates decrease in larger ecosystems, ultimately resulting in positively-sloped SARs (3). Recently, researchers have developed a trophic theory of island biogeography that more accurately predicts empirically-observed SARs, and that further generally predicts a positively-sloped relationship between food-web complexity and ecosystem size (network–area relationships, NARs) (4–8). In line with theoretical predictions, larger ecosystems are often observed to support more complex food webs including longer food chains and more trophic links per species (4–6, 8–10).

The positive SAR came to be considered as “ecology’s most general pattern” (11) and supports many biodiversity conservation or restoration plans, which often favor the protection of larger areas (12). However, there is considerable variability in the slope of SARs, which often approaches zero and is sometimes negative (1, 11, 13). This variability largely remains unexplained (1), hence undermining our capacity to predict the consequences of habitat gain or loss on biodiversity. One way to improve our understanding of SARs slopes is to incorporate trophic interactions into the classical theory of island biogeography (6, 7). In this framework, increased predation is predicted to decrease the slope of SARs. This may be seen by examining the relationship between the slope *z* of SARs at the immigration-extinction equilibrium and extinction rates (14):

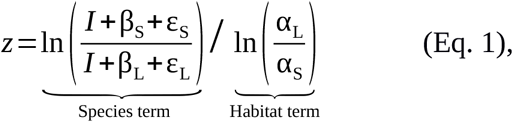

where *I* is immigration rate (assumed constant across all ecosystem sizes), and β_S_ and β_L_ are background extinction rates in a small and a large ecosystems, respectively, with β_S_>β_L_ as predicted by the theory of island biogeography. α_L_ and α_S_ are surface areas of a small and large ecosystems where predators cause additional mortality rates ε_S_ and ε_L_, respectively. The sign of the SAR slope *z* depends on the relative values of β_S_, β_L_, ε_S_ and ε_L_ .

If predator-induced extinctions ε are less sensitive to change in ecosystem size than background extinctions β, the inequality β_S_+ε_S_>β_L_+ε_L_ holds in Eq. 1, and predators flatten the SAR, but SAR’s slope *z* remains positive (14). In line with this prediction, predators were empirically shown to flatten SARs in both terrestrial and aquatic systems (14–16). In contrast, if ε scales more steeply than β with ecosystem size, the inequality flips to β_S_+ε_S_<β_L_+ε_L_, and the SAR is reversed to negative (6).

This later scenario is biologically plausible, because predators are more abundant in larger habitats (4, 6, 10, 15, 17), and more prevalent predators drive higher prey extinction rates (18). However, this scenario requires that predators are strong enough as drivers of prey extinctions and, at the same time, are generalist and omnivorous enough so that they do not suffer from decreased prey richness (6, 7). Typically, these two conditions are fulfilled by invasive alien predators (18–22), which thus have a high potential to reverse SARs to negative. Finally, because food-web properties are so strongly dependent on species richness (23), and because NARs are largely underlain by the SAR (9), we further expected a SAR reversal by invasive alien predators to also entail NARs reversal.

To empirically test these predictions, we leverage data from a long-term monitoring program in rural ponds (180 ponds, 686 sampling sessions, 5 years), where at most four different invasive alien predators may be present: the pumpkinseed fish *Lepomis gibbosus*, the black catfish *Ameiurus melas*, the mosquitofish *Gambusia affinis*, and the red swamp crayfish *Procambarus clarkii*. Freshwater ponds represent island-like habitats that are ideal to test predictions from the theory of island biogeography. In our dataset, 31 % of ponds were colonized by at least one invasive alien predator, thus providing a highly-replicated natural experimental system.

## Results

A first necessary condition for predators to be able to reverse SARs to negative is that their prevalence should increase with habitat area (6, 10). Accordingly, we found that three different prevalence metrics for invasive alien predators positively scaled with pond surface area: the probability for presence of at least one invasive alien predator, the proportion of invasive-predator richness compared to the maximum possible, and the proportional contribution of invasive predators to the local species pool (Fig. 1).

**Fig. 1.**
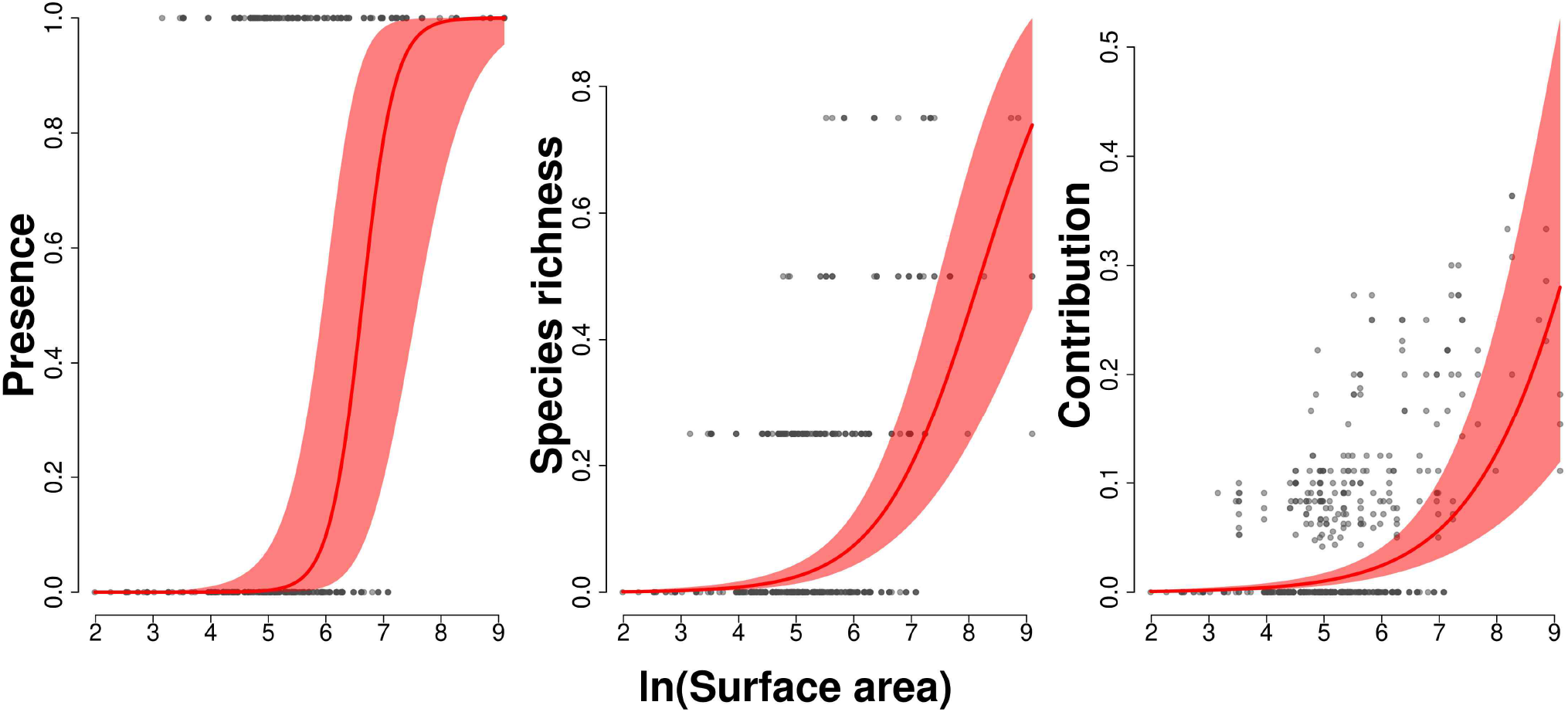
Invader prevalence-area relationships. **Left:** Probability of presence of at least one invasive predator. **Center:** Proportion of invasive-predator richness compared to the maximum possible number of invasive predators (four). **Right:** Proportional contribution of invasive species to whole-species richness. Pond surface area (m ^2^) is natural log-transformed. Grey symbols: raw data points from 686 sampling sessions performed across 180 ponds during a 5-year period. Lines and ribbons: predictions with 95 % confidence intervals from binomial generalized linear mixed models. Parameter estimates and their statistical significance for all models are provided in Table S2.

Another necessary condition for predators to be able to reverse SARs to negative is their trophic generalism, making them less sensitive to a decrease in prey richness. Measuring predator generalism required to reconstruct the “metaweb” of all trophic interactions among all species present in ponds. We reconstructed both an “observed” and an “imputed” metawebs (see Methods). In short, the observed metaweb was directly reconstructed from an extended literature survey of predator-prey relationships among all trophic species present in the ponds (825 peer-reviewed articles screened), while the “imputed” metaweb included additional trophic links that were predicted to be missing from the literature based on a modeling approach. The imputed metaweb allowed us to test for the sensitivity of our results to possible link missingness in the literature.

The observed metaweb featured 61 trophic species and 670 links (Fig. 2), while the imputed metaweb featured 846 links (Fig. S1). In both the observed and imputed metawebs, the highest-ranking predator was the grass snake *Natrix natrix*, which trophic level was 4.46 (observed) or 4.42 (imputed). The most generalist consumers were a native dragonfly-damselfly group (Aeshnidae and Libellulidae), with 30 (observed) and 39 (imputed) prey. In both metawebs, there was a positive relationship between species trophic level and trophic generalism, as measured by both indegree (prey richness) and omnivory (Figs. 3 and S2).

**Fig. 2.**
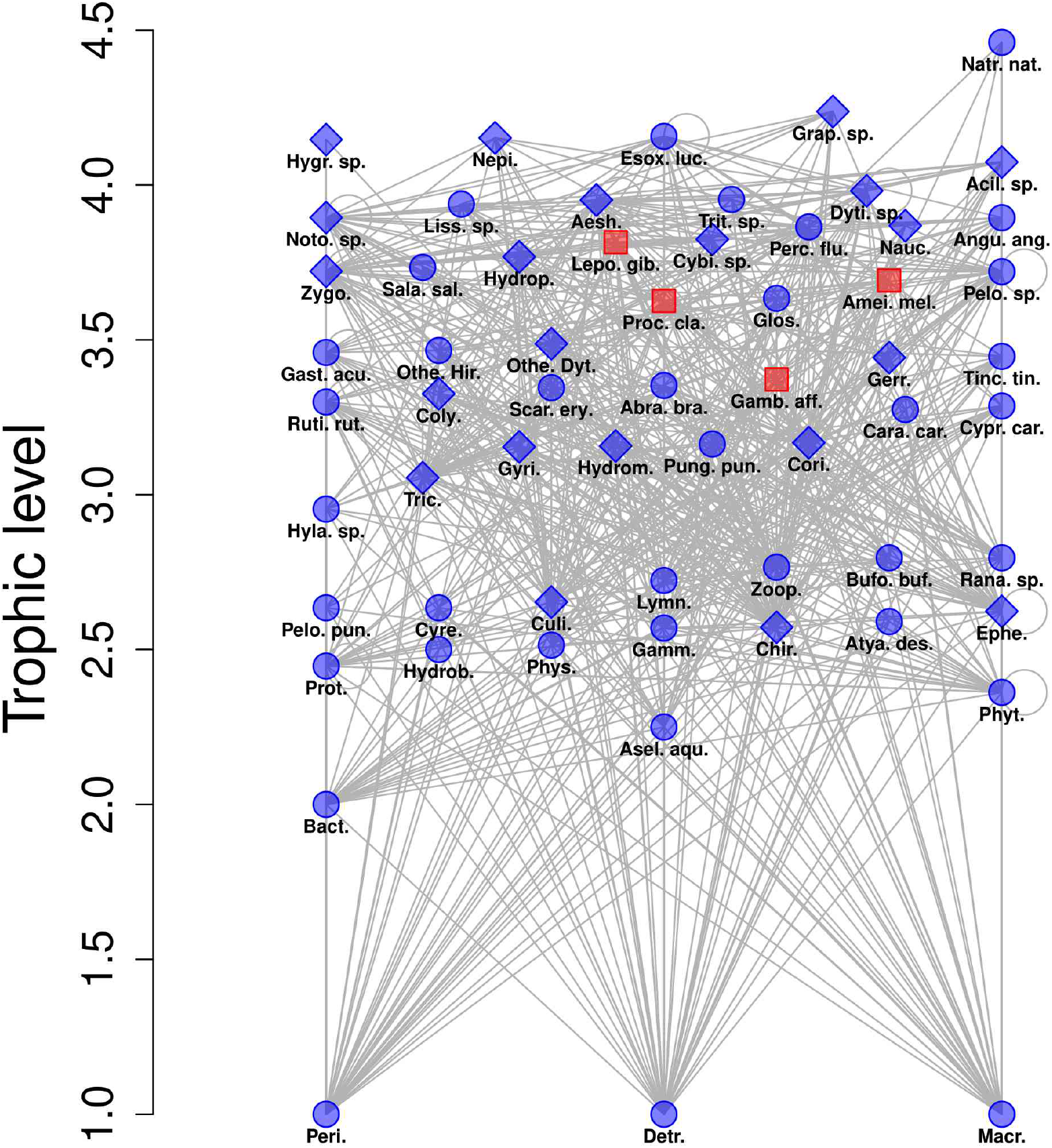
Observed trophic metaweb for Brière ponds. The reconstructed metaweb consisted of 61 trophic species (nodes) and 670 links, yielding a connectance of 0.18. Blue diamonds: native insects, which showed the strongest response to invaders (Fig. S4); Blue circles: other native species; Red squares: invaders; Lines: trophic links (directed from bottom to top, self loops are due to cannibalism). Abra. bra.: *Abramis brama*; Acil. sp.: *Acilius* sp.; Aesh.: Aeshnidae; Amei. mel.: *Ameiurus melas*; Angu. ang.: *Anguilla anguilla*; Asel. aqu.: *Asellus aquaticus*; Atya. des.: *Atyaephyra desmarestii*; Bact.: Bacteria; Bufo. buf.: *Bufo bufo*; Cara. car.: *Carassius carassius*; Chir.: Chironomidae; Coly.: Colymbetinae; Cori.: Corixidae; Culi.: Culicidae; Cybi. sp.: *Cybister* sp.; Cypr. car.: *Cyprinus carpio*; Cyre.: Cyrenidae; Detr.: Detritus; Dyti. sp.: *Dytiscus* sp.; Ephe.: Ephemeroptera; Esox. luc.: *Esox lucius*; Gamb. aff.: *Gambusia affinis*; Gamm.: *Gammaridae*; Gast. acu.: *Gasterosteus aculeatus*; Gerr.: Gerridae; Glos.: Glossiphoniidae; Grap. sp.: *Graphoderus* sp.; Gyri.: Gyrinidae; Hydrob.: Hydrobiidae; Hydrom.: Hydrometridae; Hydrop.: Hydrophilidae; Hygr. sp.: *Hygrobia* sp.; Hyla. sp.: *Hyla* sp.; Lepo. gib.: *Lepomis gibbosus*; Liss. sp.: *Lissotriton* sp.; Lymn.: *Lymnaeidae*; Macr.: Macrophytes; Natr. nat.: *Natrix natrix*; Nauc.: *Naucoridae*; Nepi.: *Nepidae*; Noto. sp.: *Notonecta* sp.; Othe. Dyt.: Other Dytiscidae; Othe. Hir.: Other Hirudinea; Pelo. pun.: *Pelodytes punctatus*; Pelo. sp.: *Pelophylax* sp.; Perc. flu.: *Perca fluviatilis*; Peri.: Periphyton; Phys.: Physidae; Phyt.: Phytoplankton; Proc. cla.: *Procambarus clarkii*; Prot.: Protozoa; Pung. pun.: *Pungitius pungitius*; Rana. sp.: *Rana* sp.; Ruti. rut.: *Rutilus rutilus*; Sala. sal.: *Salamandra salamandra*; Scar. ery.: *Scardinius erythrophthalmus*; Tinc. tin.: *Tinca tinca*; Tric.: Trichoptera; Trit. sp.: *Triturus* sp.; Zoop.: Zooplankton; Zygo.: Zygoptera. The imputed metaweb is shown in Fig. S1.

**Fig. 3.**
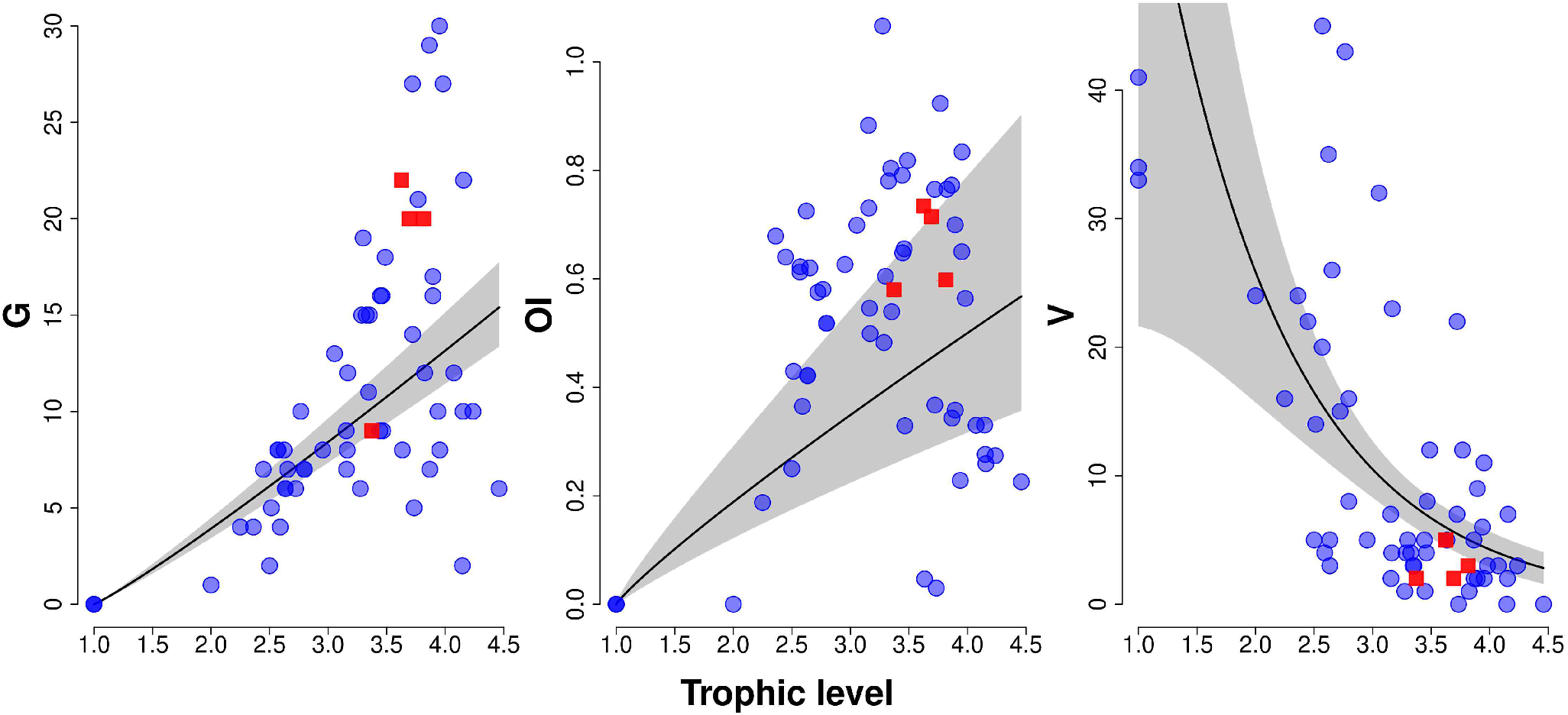
Trophic relationships in the observed metaweb. Blue circles: native trophic species, red squares: invasive alien predators; solid black lines: predicted mean relationships; shaded areas: 95% confidence intervals. Each data point represents a trophic species (n = 61). **G**: indegree (or generality) measured as the number of prey; **OI**: omnivory index computed as the weighted variance of the trophic levels of a consumer’s prey; **V**: vulnerability (or outdegree) measured as the number of predators. Predicted relationships were formed using a negative binomial GLM with log link (V), or log-log linear regressions (G, OI). Trophic relationships in the imputed metaweb are shown in Fig. S2.

Invasive alien predators were located at high trophic levels (Figs. 3 and S2), and they were hence enjoying both a high indegree and omnivory at the same time. Specifically, three of the four invasive alien predators were far above the mean indegree-trophic level relationship estimated for the 61 trophic species, indicating a disproportionately high prey richness, and all four invasive alien predators were located in the higher range of the omnivory-trophic level relationship (Fig. 3). Additionally, by occupying high trophic levels, invasive alien predators further enjoyed a relatively low vulnerability to other predators (Figs. 3 and S2), a feature theoretically-predicted to favor invasion success (22).

Finally, the last step of our analysis was to examine whether invasive alien predators were indeed able to reverse the signs of SAR and NARs slopes. At each of the 686 sampling sessions, we obtained species richness S needed to estimate the SAR by summing the total number of trophic species locally present. We further reconstructed local food webs by sampling both the observed and imputed metawebs for locally-present trophic species and their links. For each of the resultant 686 x 2 local food webs, we computed five different complexity-related network indices that, together, convey holistic information on food-web complexity (see Methods).

We explored surface-by-invader interactions on S and network indices using generalized linear mixed models (GLMMs, see Methods). This framework allowed us to accommodate the very different distributions of these variables, as well as to tease apart the confounding effects of the sampling technique, site and year. GLMMs further made it possible to account for the confounding effects of fish predators, which are known to downgrade pond biodiversity (24). This way, the estimated SARs and NARs were fish-independent and captured only the specific effects of predators being invasive aliens (for more details, see Methods).

GLMMs show that, in absence of invasive alien predators, ponds qualitatively displayed the expected positive SAR (Figs. 4 and S3). NARs were also as expected: increased species richness in larger ponds supported more complex food webs featuring more links per species (L/S), a lower connectance (C), larger mean food-chain lengths (MFCL), a higher mean trophic level (MTL), and a higher mean omnivory index of consumers (MOI). Results were very similar when local food webs were reconstructed from either the observed (Fig. 4) or imputed metawebs (Fig. S3).

**Fig. 4.**
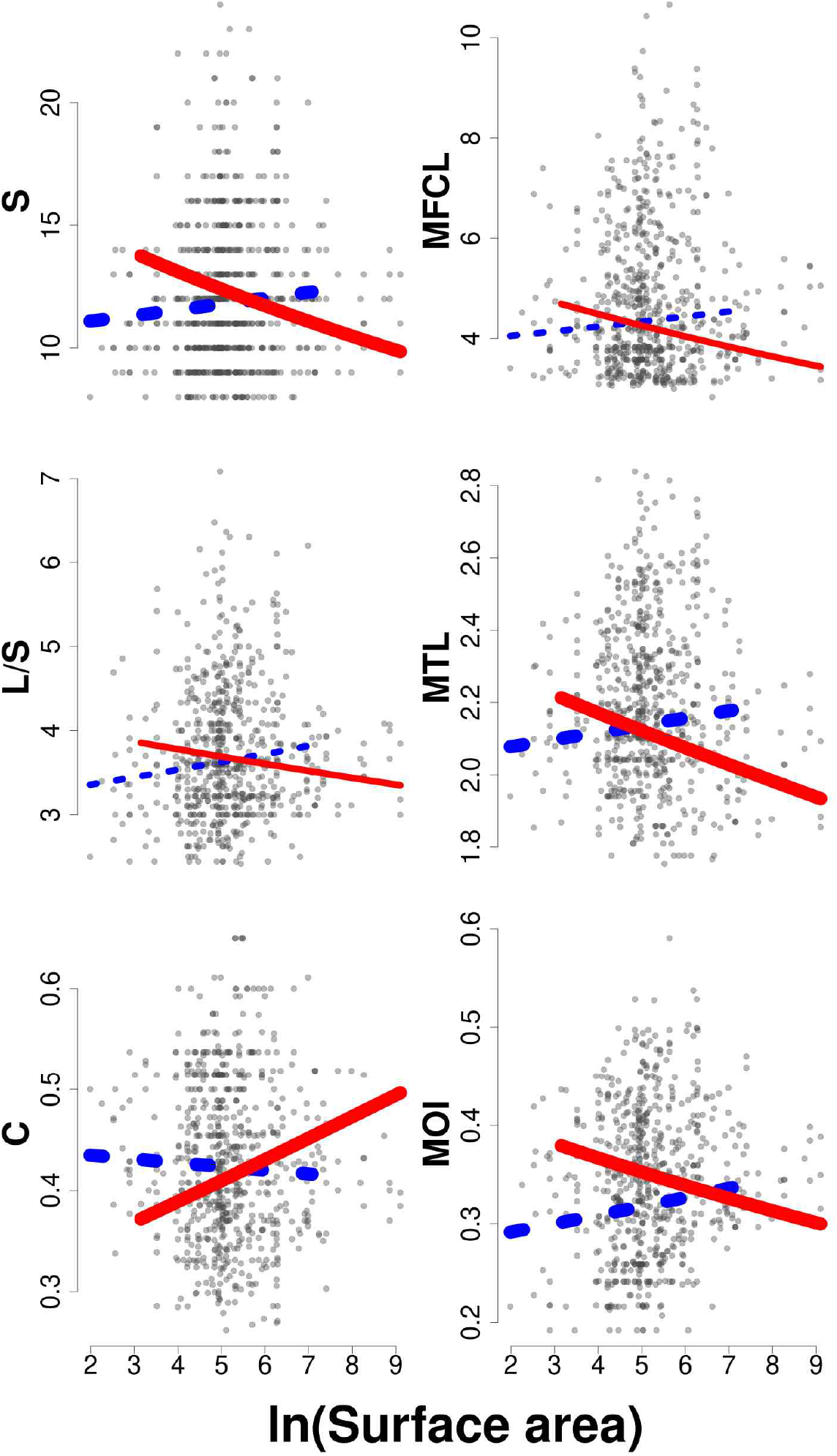
Effects of invasive alien predators on the SAR and NARs. Grey symbols: raw data; each data point corresponds to a sampling session. Lines: model predictions for a statistically-significant surface-by-invasion interaction at a 8.3 10^-3^ risk (Bonferroni correction, bold lines) or at a classical 5.0 10^-2^ risk (plain lines). Blue, dotted lines: invasive species absent; Red, solid lines: invasive species present. Predictions cover the observed range of pond surface areas for non-invaded (blue) or invaded ponds (red). S: species richness; L/S: number of links per species, C: connectance; MFCL: mean food-chain length, MTL: mean trophic level; MOI: mean omnivory index. Parameter estimates and their statistical significance for all models are provided in Table S3. The SAR and NARs from the imputed metaweb are shown in Fig. S3.

In line with the expectation that in Eq. 1 extinction rates induced by predation from invasive alien predators ε scale more steeply with ecosystem size than background extinction rates β, invasive alien predators flipped the SAR to negatively-sloped (Figs. 4 and S3). A taxon group-specific analysis shows that SAR reversal reflected a reversal in the prevalence-area relationship of insects, molluscs and non-crayfish crustaceans (Fig. S4). In contrast, in amphibians invasive alien predators decreased the intercept but not slope of the prevalence-area relationship (Fig. S4).

As also expected from the strong dependency of food-web complexity on species richness (23), SAR reversal by invasive alien predators entailed a reversal of all NARs. In invaded ponds, food-web L/S, MFCL, MTL and MOI all scaled negatively with pond area, and C scaled positively (Figs. 4 and S3), demonstrating a general food-web simplification in larger ponds.

## Discussion

The prevalence of invasive alien predators is massive and rising worldwide (25–27), and reversed SARs and NARs should therefore become increasingly common. However, the literature suggests they are not, since a visual evaluation suggests that only about 15 SARs were negatively-sloped in a meta-analysis of 794 SARs (1).

We believe that the detection of predator-induced SAR reversal has historically been hindered. First, their detection requires species to be assigned a trophic level and to fit the SAR separately to different trophic levels (14). Yet, following classical biogeography theory, the vast majority of SAR studies ignore trophic interactions. Second, even in absence of a trophic approach, a SAR reversal could still be conjectured from a hump-shaped SAR, as the SAR shifts from positive in small, predator-free ecosystems to negative in large, predator-rich ecosystems. However, detection of the hump requires both (i) the data to cover the range of ecosystem surfaces encompassing the hump and (ii) using nonlinear regression techniques such as polynomials or splines. For instance, using polynomial regressions, a predator-induced, hump-shaped SAR was observed in a study of amphibians in the Midwestern USA (28). Yet, none of the functional forms considered in SAR studies allows for a hump (29, 30), and most SAR studies are thus blind to SAR reversal in larger ecosystems.

Our results further highlight the tight intertwining of the SAR and NARs, which flipped symmetrically in response to invasive alien predators. This result suggests that the scaling of food-web properties with species richness (23) was robust both to invaders and to spatial variation, and confirms that NARs are strongly underlain by the SAR (8, 9). In turn, NARs explain why and how the SAR was predator-dependent in Brière ponds. Specifically, in absence of invasive alien predators, food-chain length increased in larger ponds, suggesting that habitat carrying capacity for predators increased with habitat area (6, 7). In parallel, food-web connectance decreased, and larger ponds had a less saturated niche space and were thus more susceptible to further invasion (31, 32). Hence, our study demonstrates that predation mediates a dynamic SAR-NAR feedback, where increased species richness in larger ecosystems favors longer food-chains and invasibility by alien predators which, in turn, increase prey extinction rates and flip the SAR and NARs to negative.

Our study also supports the theoretical prediction that, to be successful, invasive predators should enjoy both a low vulnerability to other predators and a high prey richness (22). Hence, as generalist and omnivorous consumers, invasive alien predators in Brière ponds were weakly sensitive to extinction of some of their prey and were “trophically-free” to enjoy an increasing niche space and lower demographic stochasticity in larger ecosystems, i.e., were free to display the positive SAR as predicted by the classical island biogeography theory (3), while at the same time they were shrinking the niche space and reversing the SAR in their prey (6, 14). Together, these results provide strong empirical support to the view that a trophic perspective may greatly enhance the predictive power of the theory of island biogeography (6, 7).

Finally, from an applied perspective, invasive alien predators could jeopardize area-based biodiversity conservation or restoration policies that assume purely-increasing SARs and NARs. If ecosystems already harbor invasive alien predators or are at risk of being invaded, increasing ecosystem area may benefit more to invaders than to natives, and may ultimately fail to increase biodiversity. Conversely, biodiversity loss from habitat loss could be overestimated if the relaxed prevalence and impacts of invasive alien predators are overlooked. Hence, our results are in line with the empirical observation that several small habitat patches often collectively harbor a higher biodiversity than a single large patch (12, 24). In a changing world, preserving not only the large, but also small ecosystems becomes increasingly relevant to enhancing global biodiversity conservation.

## Methods

### Study sites and biodiversity sampling

The 180 ponds were embedded in a hedgerow-landscape area of the Brière natural regional park, France (47°23’N, 02°12’W) (33–37). Brière is an area hosting a remarkable aquatic biodiversity, as testified by its classification as Ramsar and Natura 2000 sites, and includes a large (90 km^2^) marsh surrounded by a hedgerow landscape with hundreds of rural ponds. Pond surface area was measured by aerial photo-interpretation using both ArcGis 10 (Environmental Systems Research institute, Redlands, California, USA) and the BD ORTHO map of the French Institute for Geographic Information (2009 version). Pond surface area ranged from 7 to 8,951 m^2^ (mean = 364 m^2^).

Biodiversity data was produced by two research programs that sampled a multiplicity of taxa using different sampling gears and techniques (see Supplementary Methods). In total, our study included 61 “trophic species”, among which we classified as invaders three fish and one crayfish: the black catfish *Ameiurus melas*, mosquitofish *Gambusia affinis*, pumpkinseed fish *Lepomis gibbosus* and the red swamp crayfish *Procambarus clarkii*.

### Metaweb reconstruction

We reconstructed the trophic metaweb of the 61 trophic species by compiling data from the scientific literature. We scanned 825 peer-reviewed research articles documenting 16,797 proven pairwise trophic interactions. Despite this high sampling effort, 81 % of pairwise trophic interactions remained non-documented in the literature (NAs). Following standard practices, we constructed the “observed” metaweb by considering these NAs as link absence. That is, in the adjacency matrix of the observed metaweb, we ascribed *z*_*i, j*_=1 if there was at least one trophic link between prey *i* predator *j* and *z*_*i, j*_=0 otherwise.

However, there is a growing appreciation that trophic interactions are highly susceptible to undersampling due to the large amounts of time and efforts needed to document them, and that missing links in metawebs (NAs) should no longer be systematically equated to link absence, but should instead be predicted using adequate modeling techniques. We predicted missing links in the adjacency matrix of observed metaweb using the “matching-centrality” model, which demonstrated superior performance in capturing the underlying structure of food webs based on latent species traits of “matching” and “centrality” (38). We fitted the matching-centrality model to the adjacency matrix of the observed metaweb, and imputed a 1 value to 176 links that were observed as absent, but that were predicted as missing by the model (Supplementary Methods). As shown in the results section, using either the observed or imputed metawebs in our analyses did not change our results qualitatively, indicating that our conclusions are robust to possible trophic-link missingness in the literature.

### Local food-web descriptors for NARs

We sampled both the metaweb for locally-present trophic species and their links to reconstruct local food webs at each of the 686 sampling sessions, at each of which we computed five food-web descriptors (Supplementary Methods):

- (i) number of links L per trophic species (L/S), which is often taken as a measure of whole food-web complexity (9, 23);
- (ii) connectance (C), the scaled link density, which informs about how much of the trophic-niche space is saturated. More connected food webs are more robust to species loss or to introduction of invasive species (31, 32, 39), and more species-rich food webs are often observed to be less connected empirically (23).
- (iii) Mean food-chain length (MFCL), which is maybe the first NAR that was studied empirically. MFCL scales positively with surface area in lakes (10), presumably because higher trophic-level species need more space to form populations large enough to avoid stochastic extinctions (4, 6).
- (iv) mean trophic level (MTL) complements MFCL but, unlike MFCL, is mathematically independent from S. Food webs with larger MFCL and MTL have increased vertical diversity and longer energy channels, and are less robust to perturbations because of the higher sensitivity of top trophic levels to variations in primary production, mortality or climate warming (10, 40, 41).
- (v) Finally, mean omnivory index MOI is also independent of S. It measures the propensity of consumers to feed at more than one trophic level and is also an important determinant of food-web response to perturbations (42). In particular, food webs with more omnivorous consumers are less prone to exhibit a trophic cascade in response to perturbations to top predators (43).

### Statistical analyses

We modeled the three different invader prevalence-area relationships using standard binomial generalized linear mixed models (GLMMs):

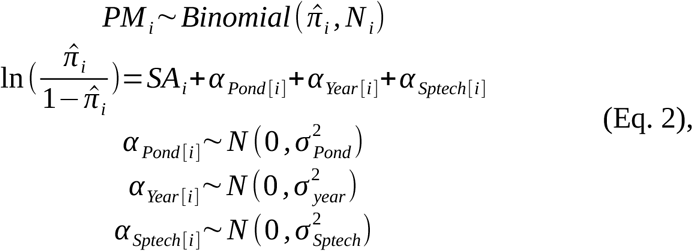

where *PM* was one of the three different prevalence metrics, *i* indexes the sampling session (n = 686), *N* was equal to one when *PM* was a 0/1 presence (hence the distribution was effectively a Bernoulli), equal to four when *PM* was invader species richness, or equal to the total number of species present when modeling prevalence as the proportion of invader species among whole-species richness (three separate GLMMs in total). *SA* was pond surface area (m^2^, natural log-transformed), standardized to zero mean and unity standard deviation. Finally, *α* _*Pond* [*i*]_, *α*_*Year* [*i*]_ and *α* _*Sptech* [*i*]_ were normally-distributed random effects of the sampling site (n = 180), year (n = 5) and technique (n = 3) on model intercept. These random effects ensured that our models accounted for all sources of non-focal variation in our data, and thus that our conclusions were immunized against false-positive errors.

We explored how S and other food-web descriptors responded to pond surface area *SA* (log-transformed) and to the presence of invasive species *Inva* using standard GLMMs:

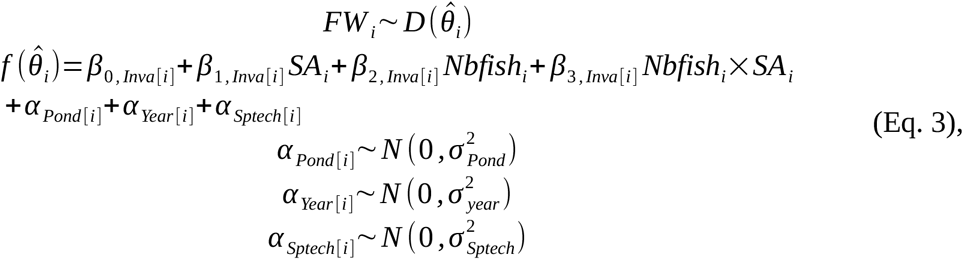

where *FW* is a food-web descriptor, *D*, 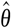 and *f* are a distribution, its parameter(s) and link function, respectively, that were specific to *FW* and selected so as to minimize model AIC. *Nbfish* is the number of fish species locally present, including both native and invasive fish species. To ease model convergence and to avoid slope-intercept correlations, all predictor variables in Eq. 3 were standardized to zero mean and unity standard deviation. Other subscripts, variables and parameters are as in Eq. 2.

In Eq. 3, the *Nbfish* effect made modeled SAR and NARs fish-independent. This independence was desirable, because fish are well-known to downgrade pond biodiversity (24). Yet, most invaders in Brière ponds were fish, and we did not want invader effects on modeled SAR and NARs to spuriously reflect fish effects. This procedure potentially generated some level of collinearity among the *Nbfish* and *Inva* predictors (¾ of invaders were fish), with the only consequence of increasing confidence intervals on their regressor estimates (44), which is a conservative effect regarding the significance of the *Inva* by *SA* interactions (i.e., collinearity protected us against false-positive errors).

A specific aspect of the SAR model (i.e., the model in which *FW* =S in Eq. 3) is that both the *Nbfish* and *Inva* effects were included in the calculation of S, which made predictors positively correlated with response by construction. However, this was unproblematic, because statistical inference was on the *Inva* by *SA* interaction only. Note also that including or excluding the *Nbfish* effect from the SAR model did not change the conclusions regarding the *Inva* by *SA* interaction on S (results not shown).

All models described in Eqs. 2 and 3 were fitted using maximum likelihood via Template Model Builder in the `glmmTMB` library of R version 4.4.2 (45, 46). Statistical significance of predictor terms in each model was tested using a Wald chi-square test in a type III analysis of deviance of each model, as provided by the `car` package in R (47). Parameter estimates from the GLMMs described in Eqs. 2 and 3 are provided in Tables S2 and S3, respectively.

## Acknowledgments

We thank the regional natural park of Brière for logistic support, the landowners for allowing us to sample their ponds, and A. Tréguier, N. Belouard, G. Surzur, A. Oger, and J.-P. Damien for assistance during fieldwork. We are grateful to Lucie Baude, Ornella Bertin and Jérémie Bruset for their contribution to compiling the literature data. All applicable ethical guidelines for the use of vertebrates in scientific research have been followed as stipulated in the licenses delivered by local authorities (license numbers 45/2011, 15/2012 and 10/2015 of the Prefecture de la Loire-Atlantique). The data used in this study were produced by previous projects funded by the Office Français de la Biodiversité (grants A10 and 221 to JMP) and by the Ministère de l’Éducation Nationale, de l’Enseignement Supérieur et de la Recherche (PhD grants to A. Tréguier and N. Belouard (JMP). No artificial intelligence was used at any stage in the production of this work.

## Supporting Information

### Supplementary Methods

Biodiversity sampling

Matching centrality model and Table S1

Computation of food-web descriptors

### Supplementary Results

Table S2. Parameter estimates for invader-prevalence relationships

Fig. S1. Imputed metaweb

Fig. S2. Trophic relationships in the imputed metaweb

Fig. S3. SAR and NARs from the imputed metaweb

Table S3. Parameter estimates for models of the SAR and NARs

Fig. S4. Effects of invasive alien predators on the prevalence of specific taxon groups

## Supplementary Methods

### Biodiversity sampling

We took advantage of the data produced by two previous research programs that sampled a diversity of taxa in Brière Ponds. The first research program (2010-2012) aimed at exploring pond invasibility patterns by the red swamp crayfish, as well as trophic constraints on crayfish invasion success (1, 2). The second program (2016-2017) aimed at exploring crayfish-amphibian-whole community interactions in ponds (3, 4). Together, these programs used multiple sampling gears:

- baited or non-baited funnel traps: 50 cm × 29 cm x 21 cm, 5.5 mm mesh size, with two entrances; and 55 cm x 17 cm x 17 cm, 5.5 mm mesh size, one entrance; set at 10-m intervals along the shoreline, in the limit of 12 per pond and laid over a day-night cycle (2).
- pipe, which consists of quickly plunging a 0.25 m^2^ hollow cylinder through the water column into the sediments, and fishing all organisms with a kick net (4 mm mesh size) until depletion; one pipe every 10 m along the shoreline in the limits of 10 samples per pond (4).
- Ortmann’s traps: empty 10-L buckets with two closed 0.33 L plastic bottles at the upper part of the bucket wall acting as floats and four openings (half-cut inverted 1.5 L plastic bottles inserted laterally (three) or at the bottom (one) acting as funnels) focusing on amphibians but also capturing by-catch macroinvertebrates swimming in the water column. Traps were set overnight approximately every 5 m along the shoreline.
- kick net (4 mm mesh size) used along the shoreline to complement sampling when specifically investigating stable isotope niches of focal species.

In total, we performed 686 sampling sessions in 180 ponds. The four different sampling gears were used either alone or in combination, thus defining three “sampling techniques”: 488 funnel-trap-only, 188 pipe-only and 10 sessions combining funnel traps, Ortmman’s traps and kick nets.

All captured organisms were determined in the field to the lowest-possible taxonomic level. When only amphibian adults were captured in ponds, we assumed juvenile presence also and we included both stages in metaweb reconstruction (see below). In total, 54 taxa were sampled, among which four were classified as invasive according to the French Resource Center on Invasive Exotic Species^1^: three fish species *Gambusia affinis, Lepomis gibbosus, Ameiurus melas* and the red swamp crayfish *Procambarus clarkii*.

We further added to the taxa list seven “trophic species”: zooplankton, phytoplankton, periphyton, macrophytes, detritus and organic matter, protozoa and protista, and bacteria. We did not sample these trophic species, but we assumed they were always present at all sampling sessions. Such an addition of ubiquitous basal resources to a sampled pool of trophic species is a common approach to represent missing trophic information when food-web samples are incomplete (5–11).

### Matching centrality model and Table S1

We inferred missing trophic links from the literature data using the matching-centrality model (12):

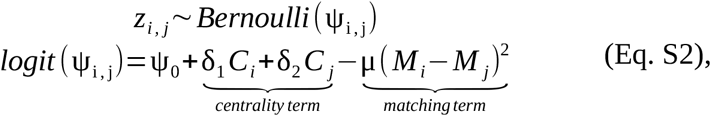

where *z*_*i, j*_ is the entry for prey *i* (row) and predator *j* (column) in the observed adjacency matrix of the metaweb, as recorded from the literature search (*z*_*i, j*_=1 if a link was present or *z*_*i, j*_=0 otherwise), *C*_*i*_, *C* _*j*_, *M*_*i*_ and *M* _*j*_ are latent traits of “centrality” and “matching”, respectively, and δ_1_, δ_2_ and μ are positive constants that scale the relative importance of prey centrality, predator centrality and predator-prey matching, respectively. The matching-centrality model was shown to outperform other modeling methods in capturing the hidden structure of interaction networks, including food webs (12).

We fitted the matching-centrality model using Markov chain Monte Carlo (MCMC) and weakly informative priors in JAGS 4.3.1 (13). We ran three chains during 45,000 iterations with a burnin phase of 40,000 iterations, and we assessed model convergence using using the Gelman–Rubin statistic (14). We further assessed goodness of fit using a Bayesian P-value (15): we computed model residuals for the actual data as well as for synthetic data simulated from estimated model parameters (i.e., residuals from fitting the model to ‘‘ideal’’ data); the Bayesian P-value is the proportion of simulations in which ideal residuals are larger than true residuals. Bayesian P-value was 0.47, indicating a very good fit.

**Table S1.** Posterior summary for main parameters in in the matching-centrality model described by Eq. S2. Rhat values inferior to 1.1 indicate a good convergence of the three MCMC chains.

| Parameter | mean | sd | Rhat |
| --- | --- | --- | --- |
| $\psi_0$ | -0.981 | 0.344 | 1.013 |
| $\delta_1$ | 1.444 | 0.163 | 1.050 |
| $\delta_2$ | 0.833 | 0.133 | 1.002 |
| $\mu$ | 0.790 | 0.146 | 1.024 |

Posterior estimates for main model parameters are provided in Table S1, and show that predator-prey links in the Brière metaweb were mostly dependent upon prey centrality, followed by predator centrality, closely followed by predator-prey matching (i.e., δ_1_>δ_2_≥μ). Mean posteriors for predicted link probabilities ψ_i, j_ ranged from 0.000 to 0.995 (0.172 ± 0.269 mean ± SD). To build the predicted adjacency matrix, we followed the procedure described by Rohr et al. (12). We retained a link as effectively predicted only when its mean posterior probability was larger than a threshold probability, defined as the probability at which the proportion of false positives in the predicted matrix (1 predicted and 0 observed) was equal to the proportion of false negatives (0 predicted and 1 observed). That is, the threshold probability was the probability at which the connectance was equal in both the predicted and observed adjacency matrices. We considered that missing links were those links that were absent in the observed adjacency matrix but present in the predicted adjacency matrix after applying the threshold criterion (12), and we imputed the metaweb by assigning these missing links with a 1 value (176 imputed links). The resultant imputed metaweb featured 846 links and a connectance of 0.23 (Fig. S1 below).

### Computation of food-web descriptors

We sampled both the observed (non-imputed) and imputed metawebs for locally-present trophic species and their links to reconstruct local food webs at each of the 686 sampling sessions. On each of the 686 local food webs, we computed the following commonly-used descriptors:

- Total trophic-species richness *S*.
- Number of trophic links *L*.
- Directed connectance *C*, computed as *C* =*L*/(*SS*_*C*_), where *S*_*C*_ is the number of non-basal, consumer species (5).
- Mean food-chain length *MFCL* was computed by enumerating the number of nodes in every trophic chain in the food web and calculating the mean. We used the `TrophicChainsStats` function of the `cheddar` package (16). A larger *MFCL* indicates a higher vertical diversity.
- For each trophic species, trophic level *TL* was computed as the weighted mean of a consumer’s prey (17):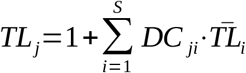, where *DC* _*ji*_ is fraction of prey *i* in the diet of the predator *j*, and 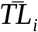 average trophic level of prey *i* . Trophic levels were computed using the `TrophInd` function of the `NetIndices` package (18). For analyses at the food-web level, we computed mean trophic level *MTL*, which reflects whether the food web web is top-or bottom-heavy.
- For each trophic species, omnivory index *OI* was computed as the weighted variance of the trophic levels of a consumer’s prey (17):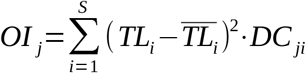, where 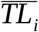 is the average of prey trophic levels. *OI*s were computed using the `TrophInd` function of the `NetIndices` package of R. For analyses at the food-web level, we computed mean omnivory index *MOI*, which reflects the mean tendency for consumers to feed at different trophic levels and is therefore a measure of vertical specialization (a lower *MOI* indicates more vertically-specialized consumers).

## Supplementary Results

**Table S2.** Parameter estimates for invader-prevalence relationships. Prevalence relationships consider the effects of habitat area on: “Presence” as the probability that at least one invasive alien predator was present in the sampling session, “Species richness” expressed as the proportion of invaders present in the sampling session relative to the maximum possible (four), and “Contribution” as the proportion of invaders among the whole-species richness. Pond surface area *SA* was standardized to unity standard deviation. sd_* effects correspond to the standard deviation of a normally-distributed random-intercept term where “pond” is pond identity, “year” is year of sampling and “sptech” is the sampling technique.

| Abundance metric | Effect | Estimate | SE of estimate | Z-value | P-value |
| --- | --- | --- | --- | --- | --- |
| Presence | (Intercept) | -5.078 | 1.540 | -3.297 | 0.0010 |
|  | SA | 3.644 | 0.955 | 3.816 | 0.0001 |
|  | sd_pond | 6.162 |  |  |  |
|  | sd_year | 0.436 |  |  |  |
|  | sd_sptech | 0.867 |  |  |  |
| Species richness | (Intercept) | -3.466 | 0.306 | -11.321 | 0.0000 |
|  | SA | 1.189 | 0.169 | 7.055 | 0.0000 |
|  | sd_pond | 1.771 |  |  |  |
|  | sd_year | 0.000 |  |  |  |
|  | sd_sptech | 0.137 |  |  |  |
| Proportion | (Intercept) | -4.408 | 0.275 | -16.022 | 0.0000 |
|  | SA | 0.914 | 0.137 | 6.686 | 0.0000 |
|  | sd_pond | 1.449 |  |  |  |
|  | sd_year | 0.054 |  |  |  |
|  | sd_sptech | 0.080 |  |  |  |

**Fig. S1.**
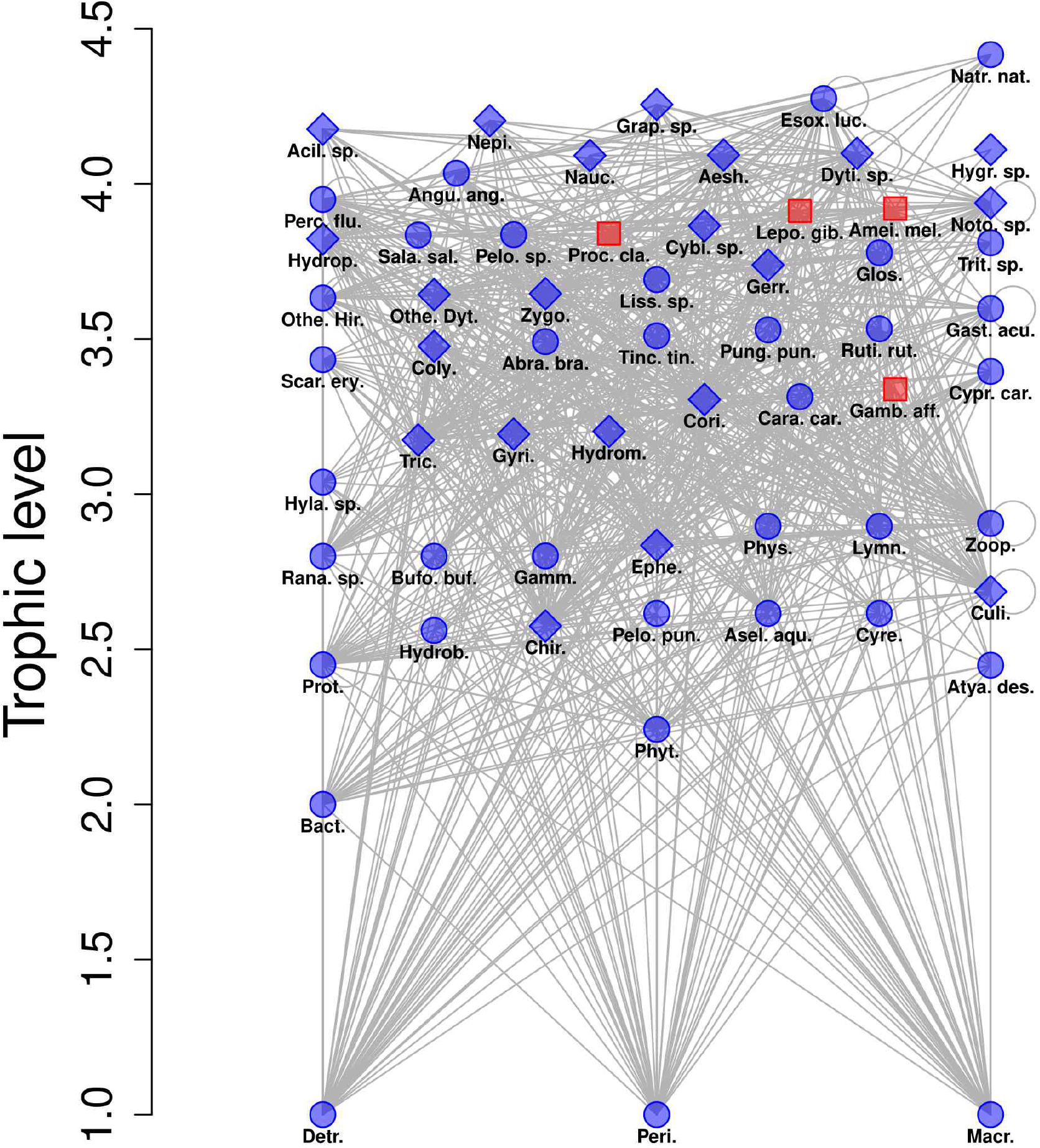
Imputed metaweb featuring 61 trophic species (nodes) and 846 links (including 176 imputed links), yielding a connectance of 0.23. Blue diamonds: insects; Blue circles: other native species; Red squares: invaders; Lines: trophic links (directed from bottom to top, self loops are due to cannibalism). Abbreviations for trophic-species names are provided in Fig. 2 caption in the main text.

**Fig. S2.**
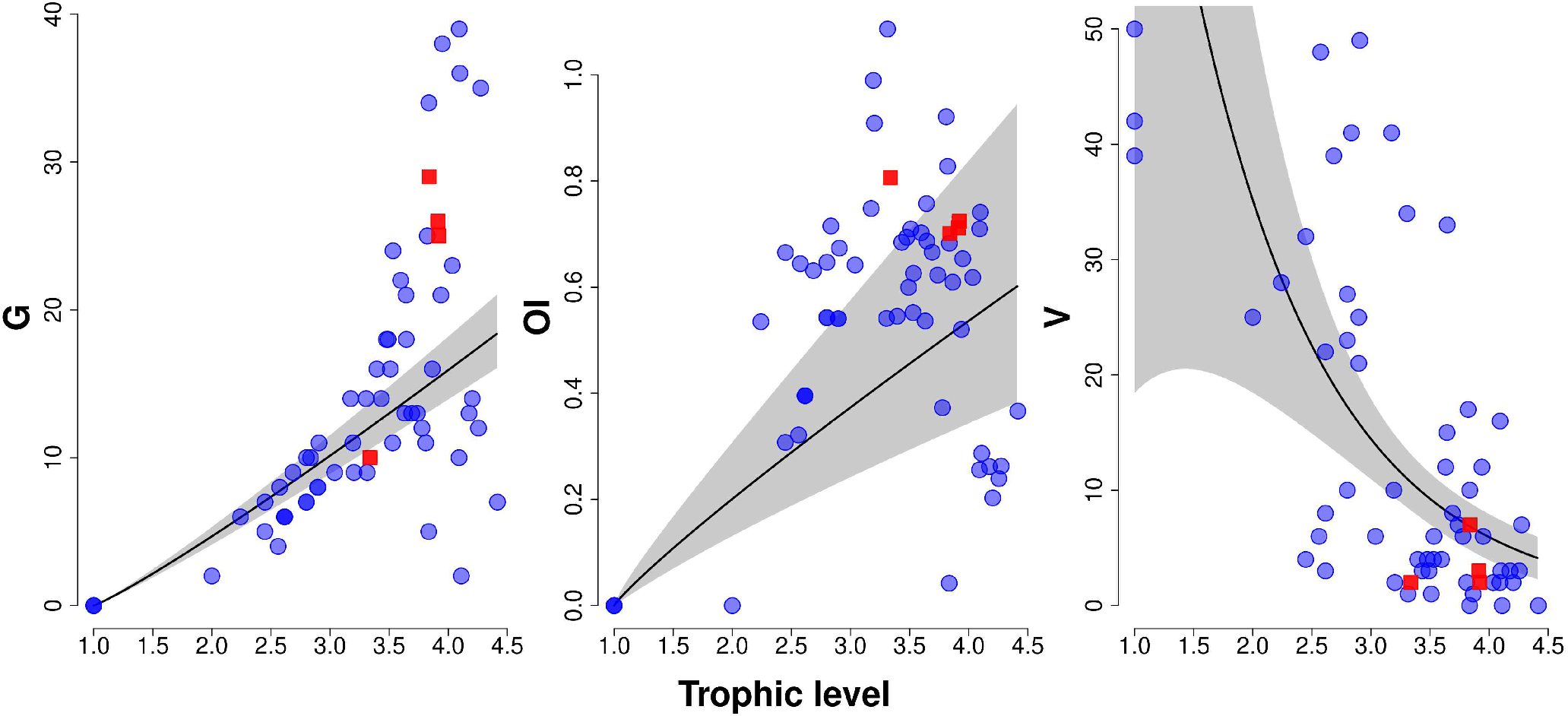
Trophic relationships in the imputed metaweb. Blue circles: native trophic species, red squares: invasive alien predators; solid black lines: predicted mean relationships; shaded areas: 95% confidence intervals. Each data point represents a trophic species (n = 61). **G**: indegree (or generality) measured as the number of prey; **OI**: omnivory index computed as the weighted variance of the trophic levels of a consumer’s prey; **V**: vulnerability (or outdegree) measured as the number of predators. Predicted relationships were formed using a negative binomial GLM with log link (V), or log-log linear regressions (G, OI).

**Fig. S3.**
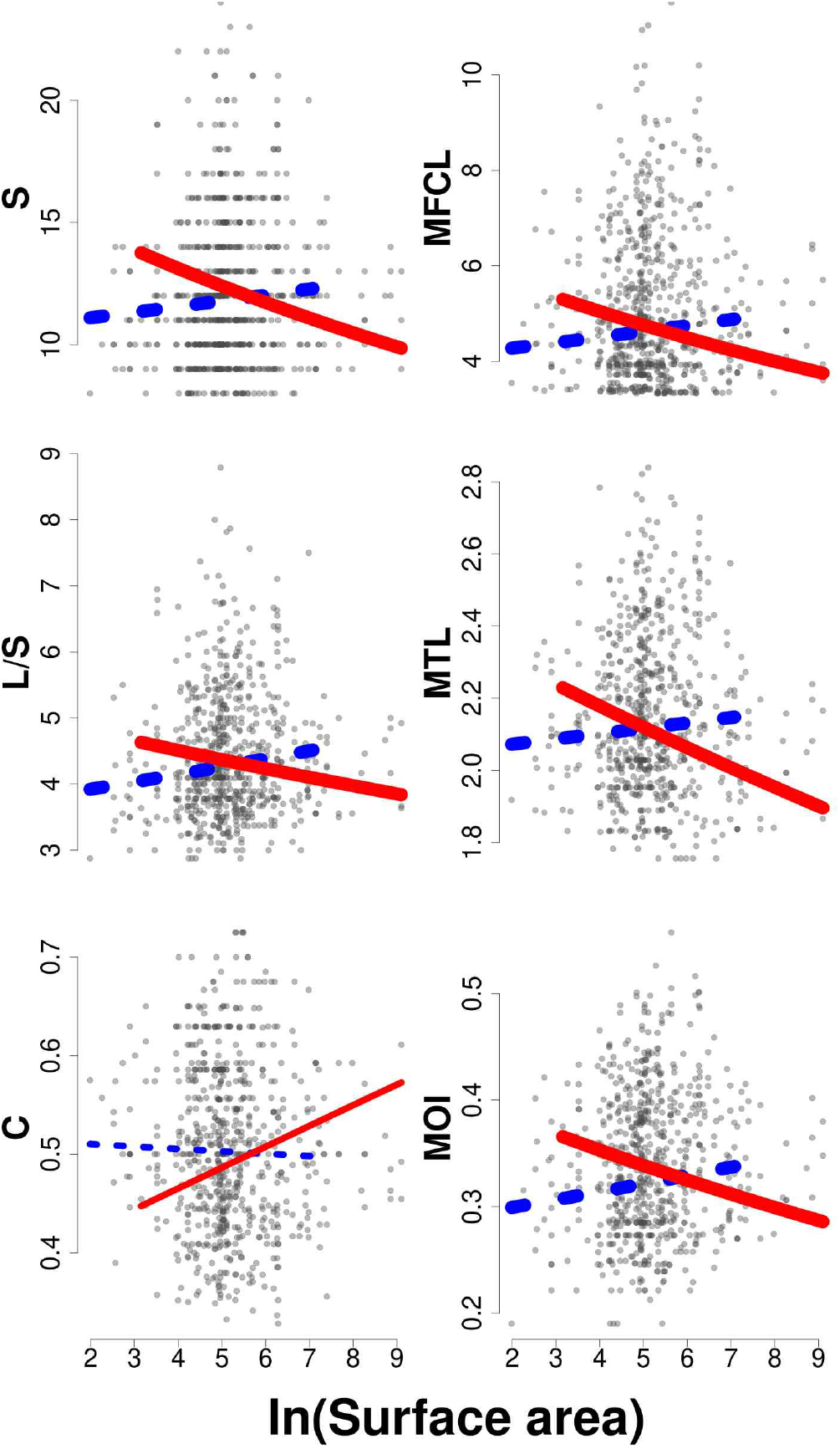
Effects of invasive alien predators on the SAR and NARs in local food webs reconstructed from the imputed metaweb. Grey symbols show the raw data (one data point per sampling session). Lines show model predictions for a statistically-significant surface-by-invasion interaction at a 8.3 10-3 risk (Bonferroni correction, bold lines) or at a classical 5.0 10-2 risk (plain lines). Blue, dotted lines: invasive species absent; Red, solid lines: invasive species present. Predictions cover the observed range of habitat surface areas for non-invaded (blue) or invaded ponds (red). S: species richness; L/S: number of links per species, C: connectance; MFCL: mean food-chain length, MTL: mean trophic level; MOI: mean omnivory index.

**Table S3.**
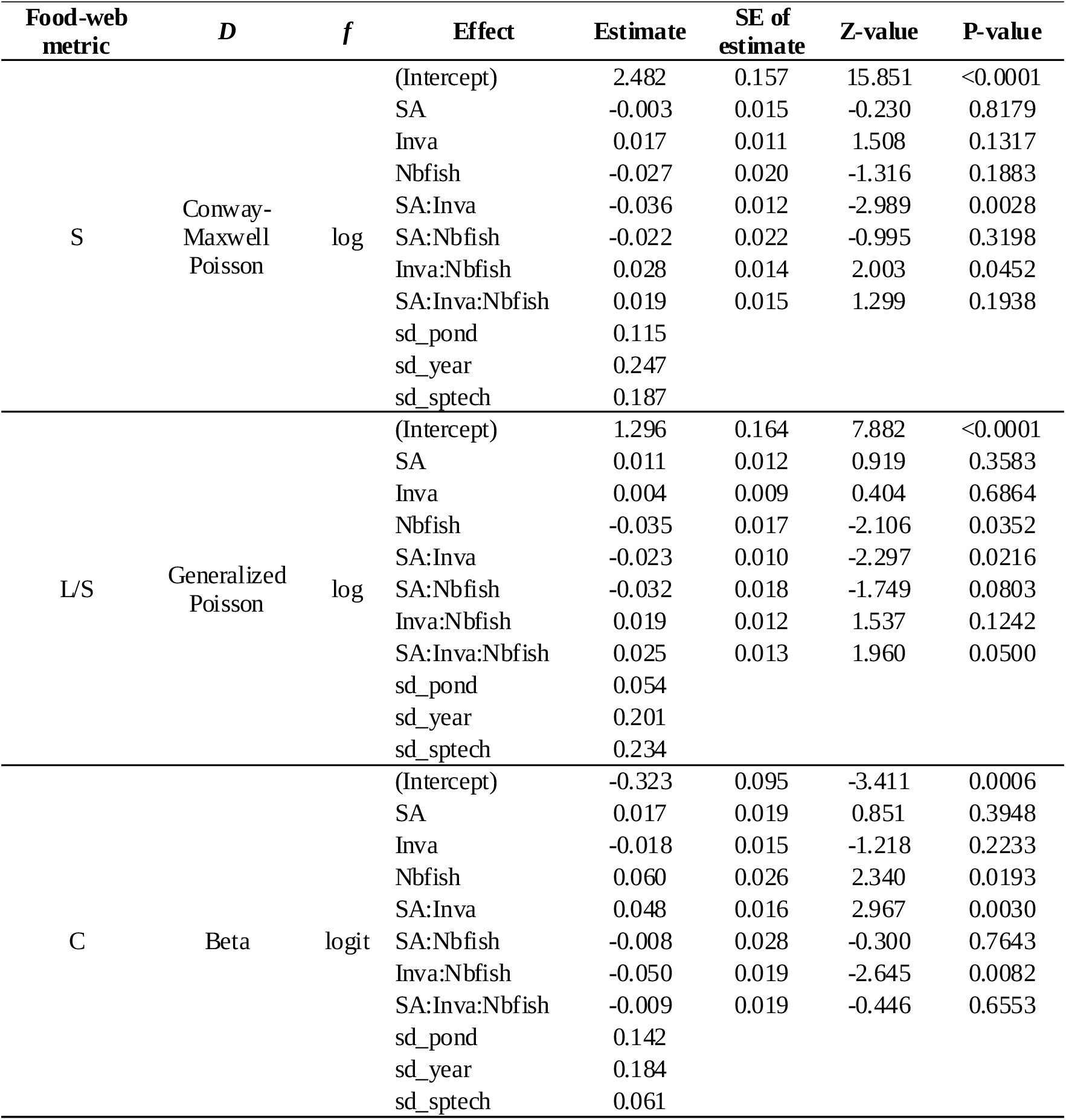
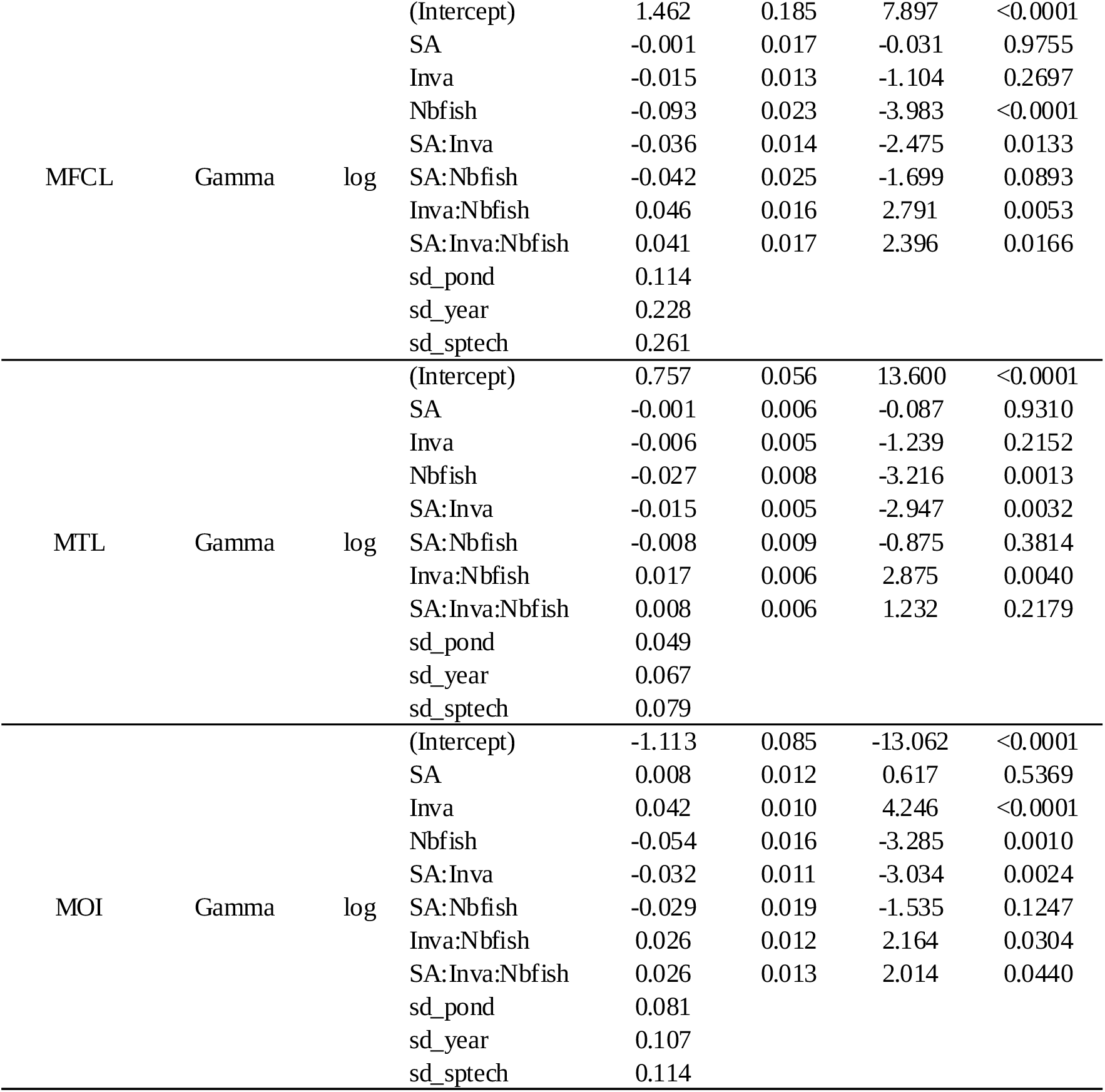
Parameter estimates for models of the SAR and NARs. Local food webs were reconstructed from the observed metaweb. Predictor variables were standardized to unity standard deviation, and estimates are thus in SD units and are interpretable as effect sizes. S: species richness; L/S: number of links per species; C: connectance; MFCL: mean food-chain length; MTL mean trophic level; MOI: mean omnivory index. sd_* effects correspond to the standard deviation of a normally-distributed random-intercept terms where “pond” is pond identity, “year” is year of sampling and “sptech” is the sampling technique (involving four different sampling gears). *D*: distribution used to model response variable. *f*: link function.

| Food-web metric | *D* | *f* | Effect | Estimate | SE of estimate | Z-value | P-value |
| --- | --- | --- | --- | --- | --- | --- | --- |
| S | Conway-Maxwell Poisson | log | (Intercept) | 2.482 | 0.157 | 15.851 | <0.0001 |
|  |  |  | SA | -0.003 | 0.015 | -0.230 | 0.8179 |
|  |  |  | Inva | 0.017 | 0.011 | 1.508 | 0.1317 |
|  |  |  | Nbfish | -0.027 | 0.020 | -1.316 | 0.1883 |
|  |  |  | SA:Inva | -0.036 | 0.012 | -2.989 | 0.0028 |
|  |  |  | SA:Nbfish | -0.022 | 0.022 | -0.995 | 0.3198 |
|  |  |  | Inva:Nbfish | 0.028 | 0.014 | 2.003 | 0.0452 |
|  |  |  | SA:Inva:Nbfish | 0.019 | 0.015 | 1.299 | 0.1938 |
|  |  |  | sd_pond | 0.115 |  |  |  |
|  |  |  | sd_year | 0.247 |  |  |  |
|  |  |  | sd_sptech | 0.187 |  |  |  |
| L/S | Generalized Poisson | log | (Intercept) | 1.296 | 0.164 | 7.882 | <0.0001 |
|  |  |  | SA | 0.011 | 0.012 | 0.919 | 0.3583 |
|  |  |  | Inva | 0.004 | 0.009 | 0.404 | 0.6864 |
|  |  |  | Nbfish | -0.035 | 0.017 | -2.106 | 0.0352 |
|  |  |  | SA:Inva | -0.023 | 0.010 | -2.297 | 0.0216 |
|  |  |  | SA:Nbfish | -0.032 | 0.018 | -1.749 | 0.0803 |
|  |  |  | Inva:Nbfish | 0.019 | 0.012 | 1.537 | 0.1242 |
|  |  |  | SA:Inva:Nbfish | 0.025 | 0.013 | 1.960 | 0.0500 |
|  |  |  | sd_pond | 0.054 |  |  |  |
|  |  |  | sd_year | 0.201 |  |  |  |
|  |  |  | sd_sptech | 0.234 |  |  |  |
| C | Beta | logit | (Intercept) | -0.323 | 0.095 | -3.411 | 0.0006 |
|  |  |  | SA | 0.017 | 0.019 | 0.851 | 0.3948 |
|  |  |  | Inva | -0.018 | 0.015 | -1.218 | 0.2233 |
|  |  |  | Nbfish | 0.060 | 0.026 | 2.340 | 0.0193 |
|  |  |  | SA:Inva | 0.048 | 0.016 | 2.967 | 0.0030 |
|  |  |  | SA:Nbfish | -0.008 | 0.028 | -0.300 | 0.7643 |
|  |  |  | Inva:Nbfish | -0.050 | 0.019 | -2.645 | 0.0082 |
|  |  |  | SA:Inva:Nbfish | -0.009 | 0.019 | -0.446 | 0.6553 |
|  |  |  | sd_pond | 0.142 |  |  |  |
|  |  |  | sd_year | 0.184 |  |  |  |
|  |  |  | sd_sptech | 0.061 |  |  |  |
| MFCL | Gamma | log | (Intercept) | 1.462 | 0.185 | 7.897 | <0.0001 |
|  |  |  | SA | -0.001 | 0.017 | -0.031 | 0.9755 |
|  |  |  | Inva | -0.015 | 0.013 | -1.104 | 0.2697 |
|  |  |  | Nbfish | -0.093 | 0.023 | -3.983 | <0.0001 |
|  |  |  | SA:Inva | -0.036 | 0.014 | -2.475 | 0.0133 |
|  |  |  | SA:Nbfish | -0.042 | 0.025 | -1.699 | 0.0893 |
|  |  |  | Inva:Nbfish | 0.046 | 0.016 | 2.791 | 0.0053 |
|  |  |  | SA:Inva:Nbfish | 0.041 | 0.017 | 2.396 | 0.0166 |
|  |  |  | sd_pond | 0.114 |  |  |  |
|  |  |  | sd_year | 0.228 |  |  |  |
|  |  |  | sd_sptech | 0.261 |  |  |  |
| MTL | Gamma | log | (Intercept) | 0.757 | 0.056 | 13.600 | <0.0001 |
|  |  |  | SA | -0.001 | 0.006 | -0.087 | 0.9310 |
|  |  |  | Inva | -0.006 | 0.005 | -1.239 | 0.2152 |
|  |  |  | Nbfish | -0.027 | 0.008 | -3.216 | 0.0013 |
|  |  |  | SA:Inva | -0.015 | 0.005 | -2.947 | 0.0032 |
|  |  |  | SA:Nbfish | -0.008 | 0.009 | -0.875 | 0.3814 |
|  |  |  | Inva:Nbfish | 0.017 | 0.006 | 2.875 | 0.0040 |
|  |  |  | SA:Inva:Nbfish | 0.008 | 0.006 | 1.232 | 0.2179 |
|  |  |  | sd_pond | 0.049 |  |  |  |
|  |  |  | sd_year | 0.067 |  |  |  |
|  |  |  | sd_sptech | 0.079 |  |  |  |
| MOI | Gamma | log | (Intercept) | -1.113 | 0.085 | -13.062 | <0.0001 |
|  |  |  | SA | 0.008 | 0.012 | 0.617 | 0.5369 |
|  |  |  | Inva | 0.042 | 0.010 | 4.246 | <0.0001 |
|  |  |  | Nbfish | -0.054 | 0.016 | -3.285 | 0.0010 |
|  |  |  | SA:Inva | -0.032 | 0.011 | -3.034 | 0.0024 |
|  |  |  | SA:Nbfish | -0.029 | 0.019 | -1.535 | 0.1247 |
|  |  |  | Inva:Nbfish | 0.026 | 0.012 | 2.164 | 0.0304 |
|  |  |  | SA:Inva:Nbfish | 0.026 | 0.013 | 2.014 | 0.0440 |
|  |  |  | sd_pond | 0.081 |  |  |  |
|  |  |  | sd_year | 0.107 |  |  |  |
|  |  |  | sd_sptech | 0.114 |  |  |  |

**Fig. S4.**
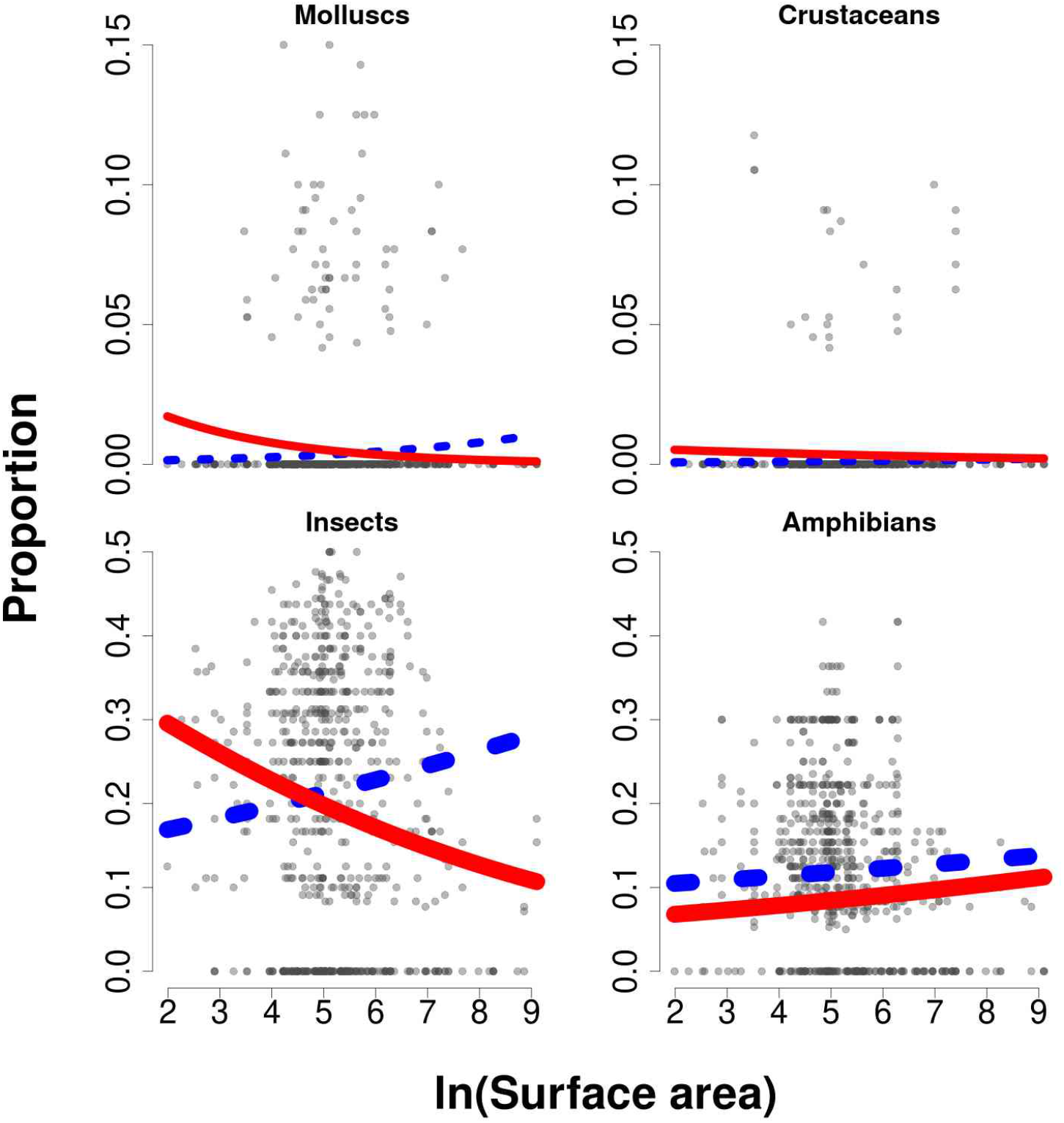
Effects of invasive alien predators on the prevalence-area relationship of specific taxon groups. Prevalence is expressed as the proportion of each taxon group to trophic-species richness at each sampling session. Grey symbols show the raw data. Lines show model predictions for a statistically-significant invasion effect or surface area-by-invasion interaction at either a 5·10^-3^ risk (thick lines) or at the standard 5·10^-2^ risk (thin lines), as tested using a Wald chi-square test in a type III analysis of deviance. Blue, dotted lines: invasive alien predators absent; Red lines: invasive alien predators present. Predictions were formed using binomial GLMMs with a linear predictor including as fixed effects pond surface area (m^2^, natural log-transformed), invader presence, and a surface-area-by-invader presence interaction, and as random-intercept effects pond identity, sampling year and sampling technique.

## Footnotes

1 https://especes-exotiques-envahissantes.fr/

